# Isolation of region-specific factors assembled on the Immunoglobulin heavy chain locus during antibody class switch recombination

**DOI:** 10.1101/2025.11.04.677290

**Authors:** Santosh Kumar Gothwal

## Abstract

Activation-Induced Cytidine Deaminase (AID) induces DNA double-strand breaks (DSBs) at the switch (S) regions of the Immunoglobulin heavy chain (*Igh*) locus, which are essential for Class Switch Recombination (CSR) and Somatic Hypermutation (SHM), key processes for effective antibody production. While AID activity is critical, its off-target effects, such as DSBs at the *Myc* locus, can cause chromosomal translocations like *Igh–Myc* fusions, contributing to B-cell lymphomas. The factors assembled on *Igh* locus that tether AID-induced DSBs and subsequently the CSR, remain unknown. To address this, we developed a method to isolate CSR-specific factors by inserting a *5X-GAL4-UAS* sequence at the switch-mu (*Sµ*) region in CH12 cells. This engineered site enables recruitment of a 3-FLAG-GAL4 DNA-binding protein (3F-GAL4-DBD), allowing specific pulldown of proteins enriched at the *Sµ* region. Successful recovery of BRD2 from the *Sµ* region, a known CSR regulator, validated this approach. Characterizing these factors may uncover novel regulators of CSR and highlight mechanisms that balance antibody diversification with genomic integrity in B cells.

## Introduction

The region-specific assembly of recombination factors is a hallmark of programmed DNA break formation across diverse recombination systems, including meiotic recombination, V(D)J recombination, and Class switch recombination (CSR). In meiotic recombination, DSBs are generated at specific genomic sites known as recombination hotspots [1-3]. The meiosis-specific topoisomerase Spo11 is recruited to these hotspots which requires prior assembly of epigenetic factors, cohesins, and the establishment of chromosomal loop-axis structures [4-7]. Similarly, V (variable), D (diversity), and J (joining) segments in immunoglobulin (*Ig*) loci undergo V(D)J recombination stimulated by RAG1/2 recombinase complex assembly on recombination signal sequences (RSS), guided by epigenetic modifications and region-specific chromosome conformation [8, 9]. In CSR, the immunoglobulin heavy chain (*Igh*) locus is proposed to harbor hotspots for AID-induced DNA breaks [10]. The RGYW motif (A/G, G, C/T, A/T), as well as short repetitive or palindromic sequences, have been identified as hot spots of somatic hypermutation and CSR {Rogozin, 1992, 1420357; [11]. These breaks occur in a DNA sequence-specific manner with chromatin conformation and epigenetic regulation playing crucial roles in determining CSR efficiency [12-14]. Although AID is essential for initiating DSB formation on the *Ig* locus, the mechanisms governing the region-specific assembly of factors that restrict AID induced DNA-breaks to the *Ig* locus while preventing off-target DNA breaks remain poorly understood.

AID activity is indispensable for initiating somatic hypermutation (SHM) and CSR, which diversify antibodies by altering the variable (V) regions to enhance antigen binding and switching antibody isotypes from IgM to IgG, IgE, or IgA [15]. However, AID targeting is not exclusively restricted to the *Ig* locus. AID can induce DNA breaks in non-*Ig* loci, such as the *Myc* and *Bcl6* loci, resulting in chromosomal translocation [15-21]. These translocations highlight dark side of AID mistargeting leading to B-lymphomagenesis. Thus, restricting AID-induced DNA breaks specifically to the *Ig* locus while limiting off-target activity is essential for ensuring the healthy humoral immune response [16, 18]. Despite AID’s discovery 24 years ago, the factors that concentrate AID activity to the *Ig* locus and those that promote off-target activity are not fully understood. It is well-documented that switch regions and non-coding RNA-driven germline transcription are critical for AID-induced DSB formation in the *Ig* locus [22]. The Sµ region plays a key role in CSR initiation, as it is occupied by specific CSR factors and forms synapses with alternative switch regions within B cells [12]. However, the factors organizing at the Sµ region simultaneously with AID expression remain unidentified

To address this question, we leveraged the *GAL4-UAS* system, originally used to regulate galactose metabolism in yeast through the binding of GAL4 transcription factor (GAL4-DBD) to the upstream activating sequence (UAS) enhancer to activate transcription [23]. By integrating a *5X-GAL4-UAS* sequence into the *Sµ* region and expressing 3F-GAL4-DBD, we developed a CH12 cell line capable of isolating CSR-specific factors by immunoprecipitation (IP) of 3F-GAL4-DBD. This modified system is proficient for AID-induced site-specific DSB formation in the flanking *Sµ* region. This study presents a novel approach for isolation of CSR factors critical for AID targeting, DNA break formation, and error-prone repair pathways. Our modified CH12 cell line serves as a powerful tool for investigating the mechanisms that establish AID hotspots, ensuring the safe and efficient execution of CSR while minimizing the risks of genomic instability and B-lymphomagenesis.

## Results

### Modification of *Igh* locus by *5X-GAL4-UAS* Insertion Upstream of the *Sµ* Region

The mouse *Ig* loci consist of coding and non-coding sequences on chromosome 12 over a distance of 324160 bp between the *Ighv5-2* and *Igha* (Gencode Transcript: ENSMUST00000194738.6). This long region harbors noncoding “switch” *S* regions at defined intervals, each upstream of a specific coding “constant” *C* gene (Figure 1A). AID-induced DSB formation takes place within switch regions, while a deletional recombination event between the two *S* regions is required for isotype switching [24]. The most upstream switch region in the *Ig* locus is *Sµ*, which is cleaved to facilitate switching from IgM to other antibody isotypes. Thus, DSB formation at *Sµ* region are essential for accepting the downstream *S* regions. Thus, *Sµ* serves as a universal donor for recombination with the downstream isotype-specific S regions.

**Figure 1.**
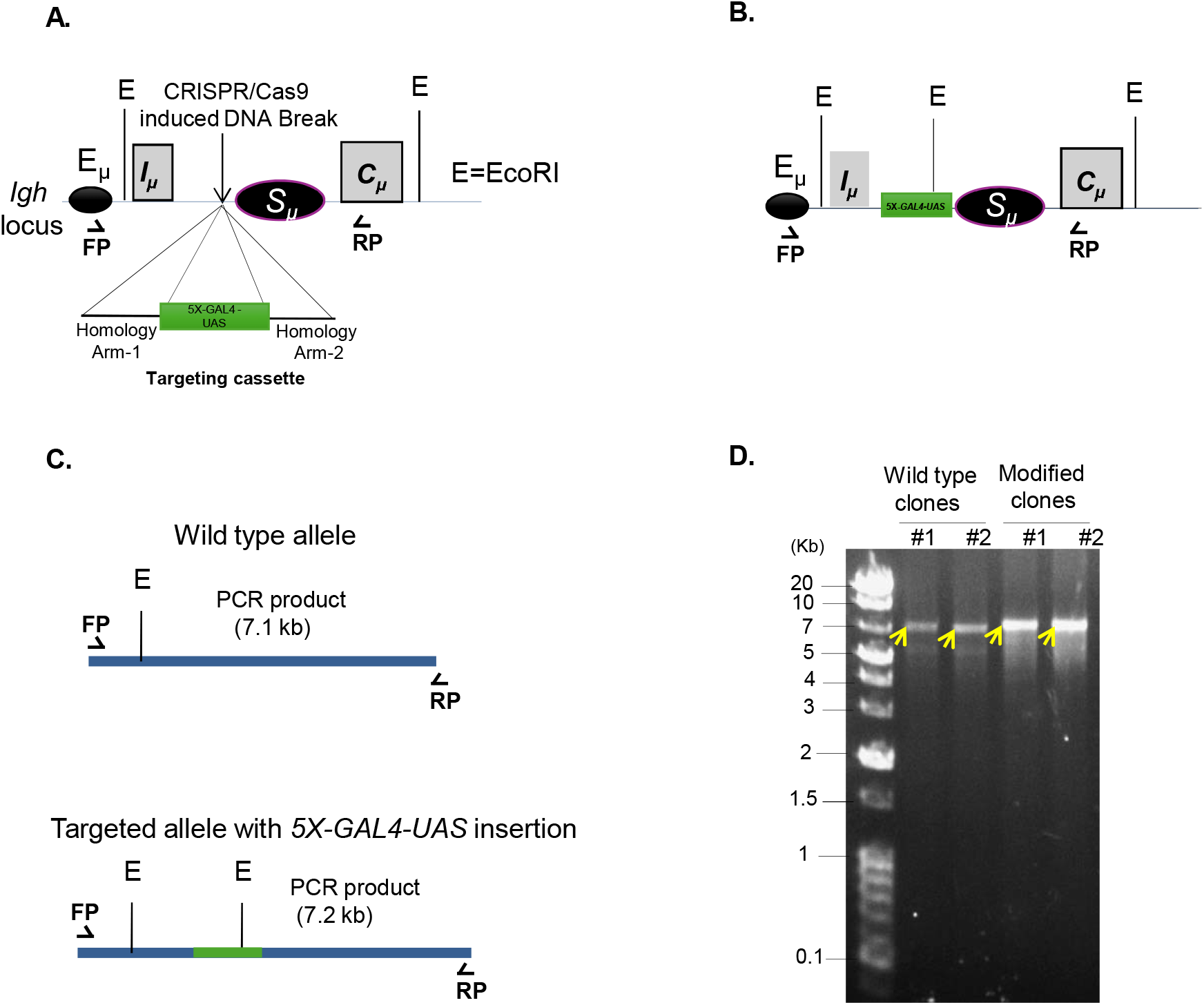
Insertion of *5X-GAL4-UAS* into the *Igh* locus of CH12 cells. **(A)** Schematic representation of *Igh* locus indicating the *Iµ* and *Sµ* location and targeting position of *5X-GAL4-UAS* sequences. *5X-GAL4-UAS* sequence flanking the homology arms upstream and downstream of CRISPR/Cas9 targeting site was used for the insertion of *5X-GAL4-UAS* into *Igh* locus. E=EcoR1 site **(B)** Schematic representation of *Igh* locus inserted with *5X-GAL4-UAS* insert upstream of *Sµ* regions. **(C)** Schematic representation of wild type and modified *Igh* locus with *5X-GAL4-UAS* insertion, indicating the primer positions. FP= Forward primer, RP= Reverse primer. **(D)** PCR amplification using 750 nanogram of genomic DNA (gDNA) from 2 independent WT clones and two independent single cell clones with the *5X-GAL4-UAS* sequence inserted are shown. The PCR product length from wild type genomic DNA using FP and RP is 7132 base pairs (bp). PCR product length using FP and RP from the *5X-GAL4-UAS* inserted clones exhibits a band of 7263 bp.

We hypothesize that *Sµ* is a critical DSB hotspot necessary for the initiation of CSR. Therefore, we inserted the *5X-GAL4-UAS* sequence, a DNA sequence from the yeast genome that can be physically bound by the GAL4 DNA-binding protein (GAL4-DBD). To avoid disrupting the assembly of CSR factors and DSB “hotspot” sequence within *Sµ*, we inserted the *5X-GAL4-UAS* approximately 1 kb upstream of *Sµ* (Figure 1A, B). This upstream insertion should allow for the natural assembly of the CSR factors that tether AID activity to the *Sµ* region (Figure 1A).

To insert *5X-GAL4-UAS*, we transfected CH12F32A cells (CH12 hereafter) with a CRISPR-Cas9 vector with a single gRNA specific to the Sµ target region (Figure 1B). CH12 cells were co-transfected with the *5X-GAL4-UAS* targeting cassette containing approximately 1 kilobase arms homologous to regions immediately upstream and downstream of the CRISPR-Cas9 target site (Figure 1A-B). After 24 hours, transfected CH12 cells were sorted for CRISPR-Cas9 plasmid expression using the co-expressed CD4 marker from plasmid backbone, followed by single-cell dilution. Pure colonies grown from single cell clones were screened for the presence of the *5X-GAL4-UAS* insertion at *Sµ* by PCR (Figure 1C-D). Since the length of the *5X-GAL4-UAS* sequence and restriction enzymes is 131 base pairs, its insertion upstream of *Sµ* region and subsequent PCR amplification with indicated primers should result in a product length of 7263 bp (Figure 1C-D). On the other hand, this PCR product length in wild-type CH12 cells gives a 7132 base pairs band due to the absence of the *5X-GAL4-UAS* and DNA-sequences of restriction enzymes (Figure 1C-D). By PCR, we confirmed a 7132 bp band in two clones of wild-type CH12 cells while two *5X-GAL4-UAS* inserted clones exhibited a 7263 bp band (Figure 1C-D). These results suggest that the clones obtained after transfection with CRISPR/Cas9 and transfection with *5X-GAL4-UAS* targeting cassette were successfully inserted with the *5X-GAL4-UAS* sequence upstream of the *Sµ* region.

### Confirmation of *5X-GAL4-UAS* Insertion into *Igh* Locus by DNA-sequencing and Southern blotting

We confirmed the integration of the *5X-GAL4-UAS* sequence in the two clones (Figure 1D). Since the PCR product length was almost indistinguishable between the wild type inserted clones due to the short length of *5X-GAL4-UAS*, we next attempted to confirm the *5X-GAL4-UAS* integration upstream of the *Sµ* region by DNA-sequencing of genomic DNA encompassing the insertion site (Figure 2A). We PCR amplified the *Igh* sequence upstream and downstream of the *5X-GAL4-UAS* insertion site and cloned the PCR product into the pZero-blunt vector followed by sequencing using forward primer located in the homology arm 1, *5X-GAL4-UAS* and the homology arm 2 (Figure 1D, Primer Table 1). Sequencing results confirmed that *Sµ* region in the arm 1 contains *5X-GAL4-UAS* sequence insertion (Figure 2A). We also confirmed the KpnI and NheI restriction sites from arm1 and arm 2, which were utilized for cloning the homology arms into the targeting cassette (Figure 2A). Off note, the KpnI sequence was point mutated, as we observed *GGTCC* instead of *GGTACC*, however, the sequence of *5X-GAL4-*UAS was fully intact without any detectable mutation while the NheI site, connecting *5X-GAL4-UAS* sequence and the homology arm 2 in the targeting construct, was present (Figure 2A). The sequencing results from gDNA of the *5X-GAL4-UAS* inserted CH12 cells confirmed that the *Sµ* region was successfully inserted with the *5X-GAL4-UAS* sequence without altering the sequences surrounding the *Sµ* region, keeping the *Ig* locus intact intact (Figure 2A).

**Table 1.**
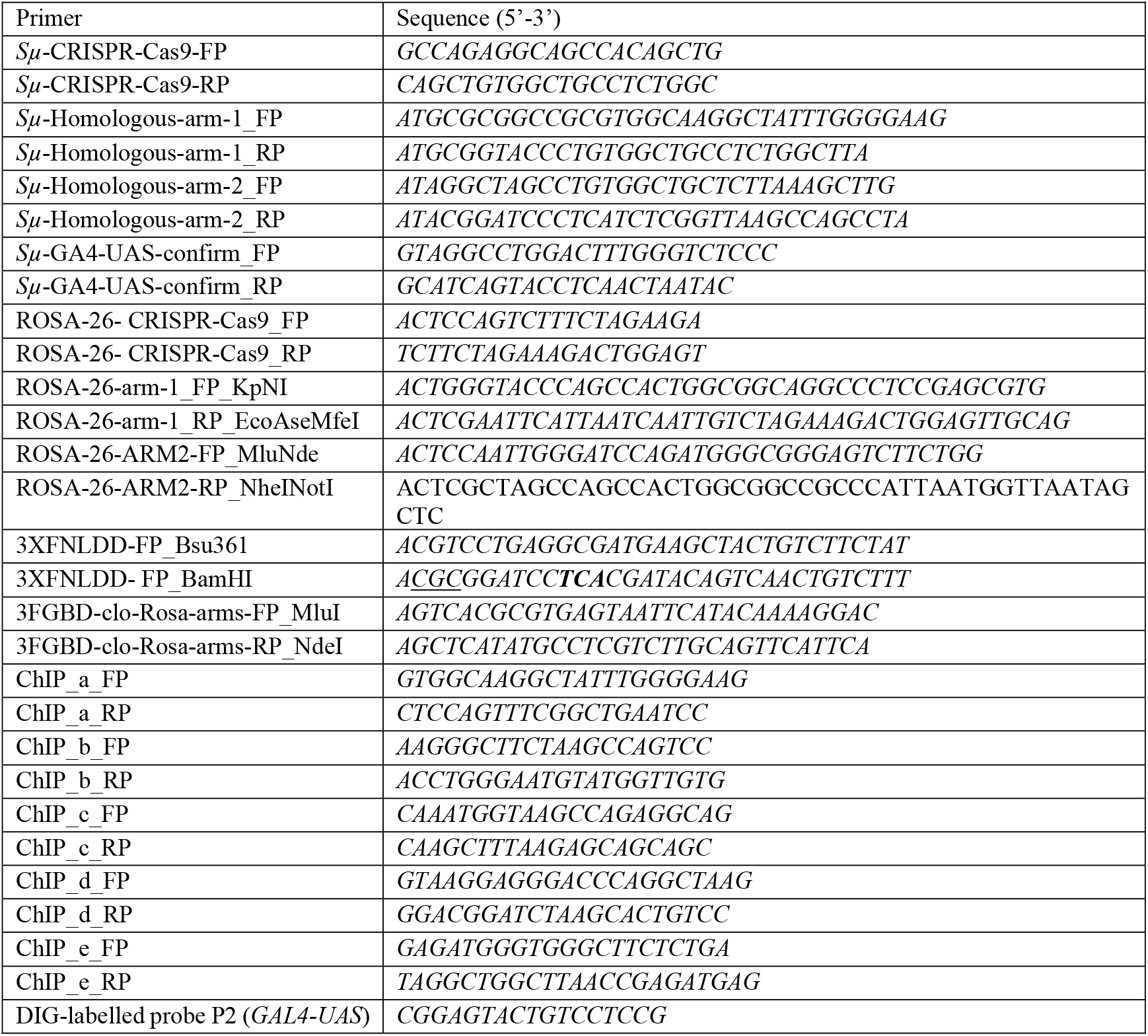
Primer sequences used in this study.

**Figure 2.**
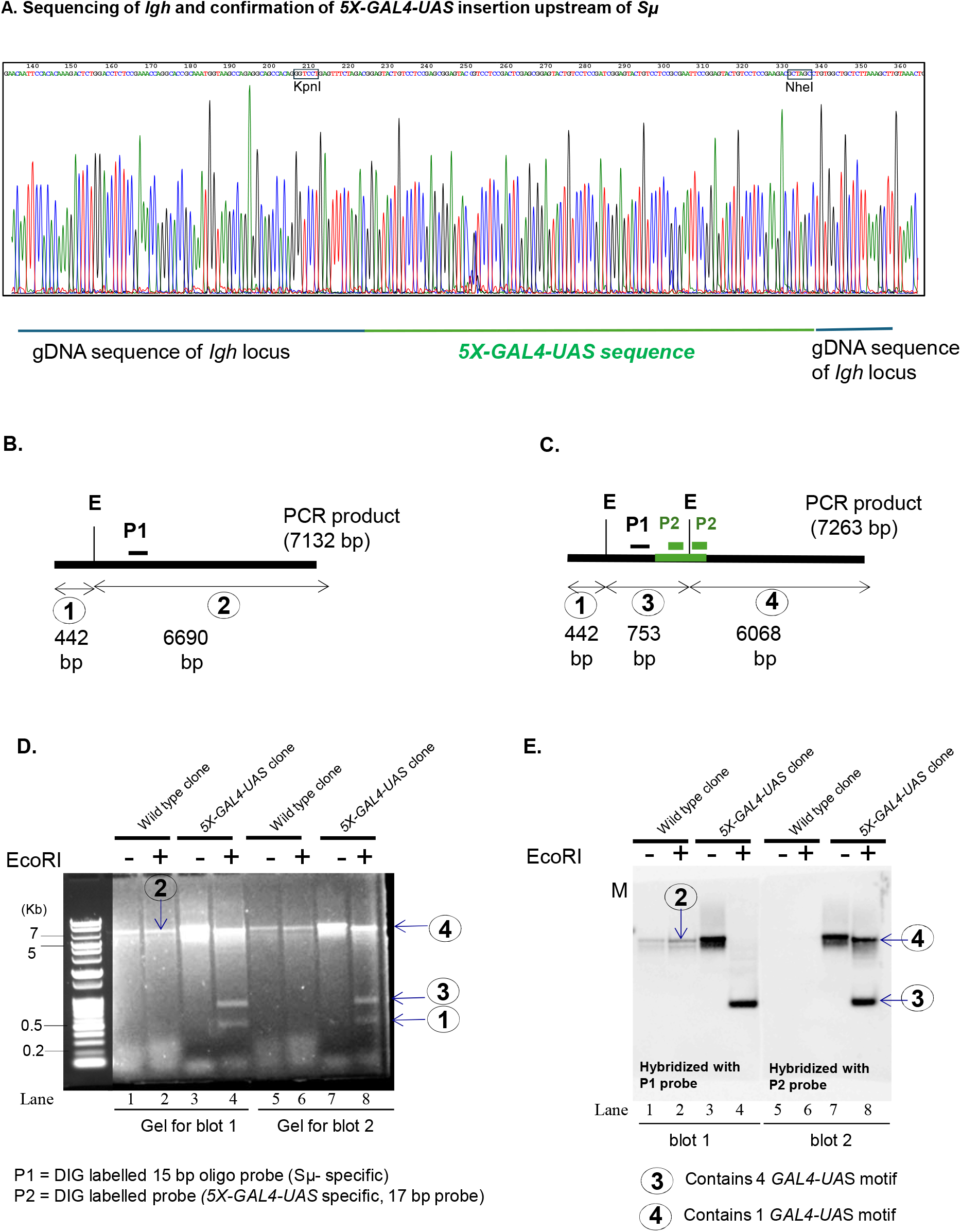
Confirmation of *5X-GAL4-UAS* insertion into the *Igh* locus of CH12 cells. **(A)** genomic DNA sequence of modified CH12 cells inserted with *5X-GA4-UAS* confirming the *5X-GA4-UAS* sequence downstream of CRISPR/Cas9 induced DSB site is shown. PCR amplifying the regions flanking the *5X-GAL4-UAS* insertion sequence was cloned in zero-blunt system (Invitrogen) and sequenced with primer, binding upstream of the *GAL4-UAS* sequence site (5’-3’ direction). The *5X-GAL4-UAS* sequence was confirmed between the sequence of homology arms 1 and 2. The KpnI and NheI restriction sites were confirmed at the junction of homology arm 1 and 2 flanking the *5X-GAL4-UAS* sequence, as constructed in the targeting cassette. **(B)** Schematic of the EcoRI digestion of PCR products obtained from wild type and *5X-GAL4-UAS* inserted clones of CH12 cells. Wild type PCR product upon the EcoRI digestion gives 2 bands (band 1 and 2) of 442 and 6690 bps. Band (1) is recognized with DIG labeled P1 probe (specific to Sµ motif) as indicated in the schematic. **(C)** *5X-GAL4-UAS* inserted cells upon PCR amplification and digestion with EcoRI exhibits three bands (band 1, 3 and 4) of 442 bp, 753 bp and 6068 bps respectively. The band (3) and band (4) harbors 4 and 1 *GAL4-UAS* motif respectively, as separated due to the EcoRI digestion. Band (3) and band (4) are recognized with P2 probe (specific to *GAL4-UAS* motif), indicated in the schematic presentation. **(D)** Ethidium bromide staining of PCR products from wild type and *5X-GA4-UAS* inserted cells with/without EcoRI digestion. The product number 1, 2, 3 and 4 were confirmed by ethidium bromide staining and indicated along with size markers in the left. Two independent clones from wild type and *5X-GAL4-UAS* inserted clones of CH12 cells were used for the confirmation of *5X-GAL-UAS* insertion by ethidium bromide staining follwed by southern hybridization with a DIG labelled probes P1 and P2. **(E)** Southern blotting and hybridization of blot 1 and blot 2 with probes P1 and P2 respectively. EcoRI-digested PCR product from wild type and *5X-GAL4-UAS* inserted clones were transferred to nitrocellulose membrane follwed by hybridization with DIG labeled probe P1 (Sµ-specific) and P2 (*GAL-UAS*-specific) respectively. The wild type PCR product upon EcoRI digestion gives 2 bands where band 2 is recognized by the P1 probe. The PCR from genomic DNA of *5X-GAL4-UAS* inserted clones gives three bands upon EcoRI digestion. Band 3 contain *4X-GAL4-UAS* motifs and Band 4 contains *1X-GAL4-UAS* motif and are recognized with the P2 probe.

We also confirmed that the full-length arms were integrated in-frame and did not alter the length of *Sµ* on the *Ig* locus. To check the full-length insertion of homology arms, we performed the PCR amplification using primers located outside the homologous recombination region of interest and further analyzed the PCR products by EcoRI digestion, follwed by ethidium bromide staining and Southern blotting with probes specific to either genomic DNA of *Igh* locus or the *5X-GAL4-UAS* regions (Figure 2B, C). EcoRI digestion of the PCR product obtained from wild type clones generates two fragments of 442 and 6690 bps (Figure 2B,D), while EcoRI digestion PCR product obtained from the *5X-GAL4-UAS* inserted clones yielded three bands of 442, 753 and 6068 bps (Figure 2C, D). The additional EcoR1 site within the *5X-GAL4-UAS* sequences led the generation of 753 and 6068 base pairs (Figure 2C, D). Additionally, Southern blotting and hybridization with probes P1 and P2 recognizing Sµ region and the *5X-GAL4-UAS* respectively, further confirmed the expected length of the DNA fragments (Figure 2E). These results confirm the *5X-GAL4-UAS* sequence was inserted in the correct frame on the *Igh* locus.

### Stable expression of 3F-GAL4-DBD in *5X-GAL4-UAS* CH12 cells

The *GAL4-UAS* sequence is recognized by the GAL4-DNA binding protein [23, 25]. Therefore, overexpression of the GAL4-DBD protein in *5X-GAL4-UAS* CH12 cells should exhibit site-specific binding to the Sµ region. We modified the GAL4-DBD by inserting a *3FLAG* sequences upstream of the GAL4-DBD protein (3F-GAL4-DBD hereafter). To efficiently purify protein complexes assembling on the *Sµ* regions, we integrated the *3F-GAL4-DBD* plasmid into the *Rosa-26* locus (Figure 3A), a transcriptionally active locus allowing the constitutive expression of inserted genes [26]. To integrate the 3F-GAL4-DBD construct into the *Rosa-26* locus, we introduced CRISPR/Cas9-induced DSBs at intron 1 followed by homology-directed replacement (Figure 3A). The *3FLAG-GAL4-DBD* is tagged with nuclear localization signals, allowing for efficient transport of 3F-GAL4-DBD to the nucleus. After transfection, single-cell dilution and puromycin selection, we selected six clones namely Cl-1, Cl-2, Cl-3, Cl-4, Cl-5, and Cl-6 and screened these for the 3F-GAL4-DBD protein expression by Western blotting analysis (Figure 3B). Among the 6 clones, Cl-2 and Cl-3 exhibited expression of 3F-GAL4-DBD protein (Figure 3B). These confirm that Cl-2 and Cl3 exhibits the *5X-GAL4-UAS* insertion and stable expression of 3F-GAL4-DBD, useful for the pulldown of CSR specific factors from the *Sµ* region (Figure 3B).

**Figure 3.**
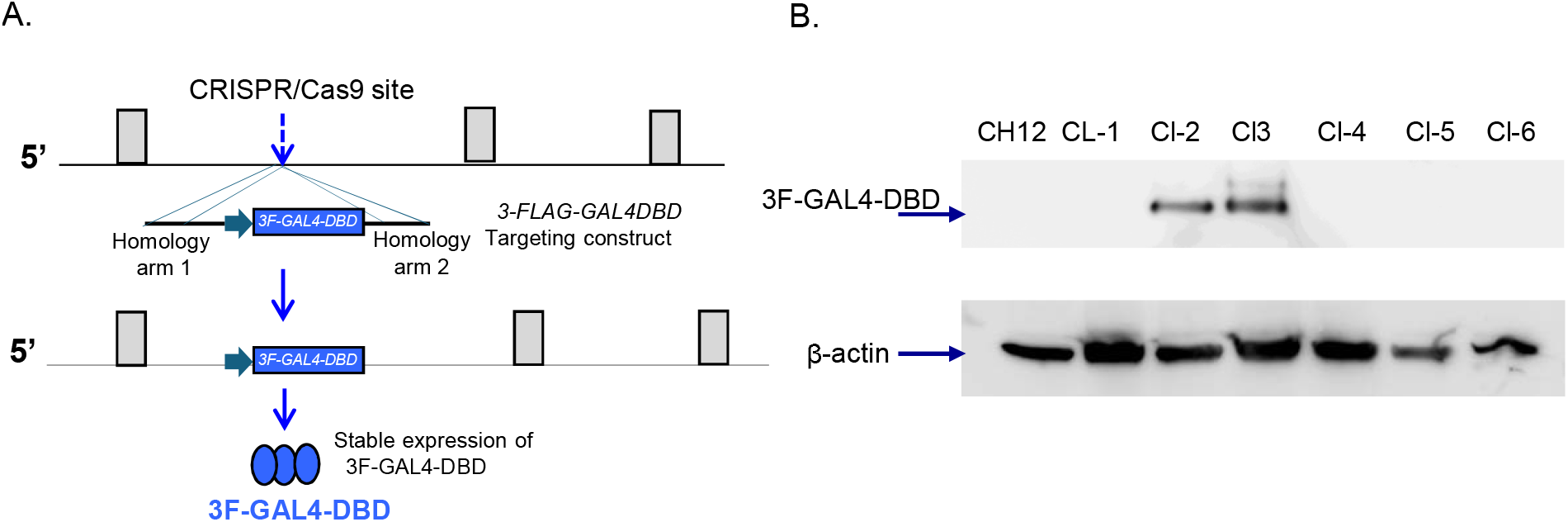
Stable expression of 3F-GAL4-DBD into the *5X-GAL4-UAS* inserted CH12 cells and confirmation of CSR induction in the modified cells. **(A)** Schematic representation of mouse *Rosa-26* locus indicating CRISPR/Cas9-induced DSB formation at intron 1 followed by replacement with the targeting construct having *pEF1-3X-Flag-GAL4-DBD-IRES-puromycin* sequences between the homology arms 1 and 2. **(B)** *5X-GAL4-UAS* inserted cells were transfected with CRISPR/Cas9 plasmid and *pEF1a-3X-Flag-GAL4-DBD-IRES-puromycin* fragment followed by single cell dilution after 24 hours of the transfection. The single cell clones were grown uptil confluency. Six clones named Cl-1, Cl-2, Cl-3, Cl-4, Cl-5 and Cl-6 were selected for the screening of 3F-GAL4-DBD expression. (B) Western blotting was performed using the whole cell lysates from Cl-1, Cl-2, Cl-3, Cl-4, Cl-5 and Cl-6 and 3F-GAL4-DBD protein expression was detected in Cl-2 and Cl-using the Flag antibody.

### Tethering of 3F-GAL4-DBD to modified *IgH* locus and CSR induction

Since the switch regions undergo chromatin conformation changes during CSR, we asked whether *5X-GAL4-UAS* clones stably expressing the 3F-GAL4-DBD undergo normal CSR, allowing the normal assembly of the CSR factor. We thus stimulated the Cl-2 and Cl-3 with CD40 ligand, IL-4 and TGF-β (CIT), inducing robust CSR to IgA in CH12 cells as described previously[13]. We noted that Cl-2 and Cl-3 exhibited 47.6% and 11.2% IgA switching, respectively suggesting that these cells undergo efficient CSR (Figure 4A). This confirms the functioning of the CSR in the *5X-GAL4-UAS* clones stably expressing the 3F-GAL4-DBD protein.

**Figure 4.**
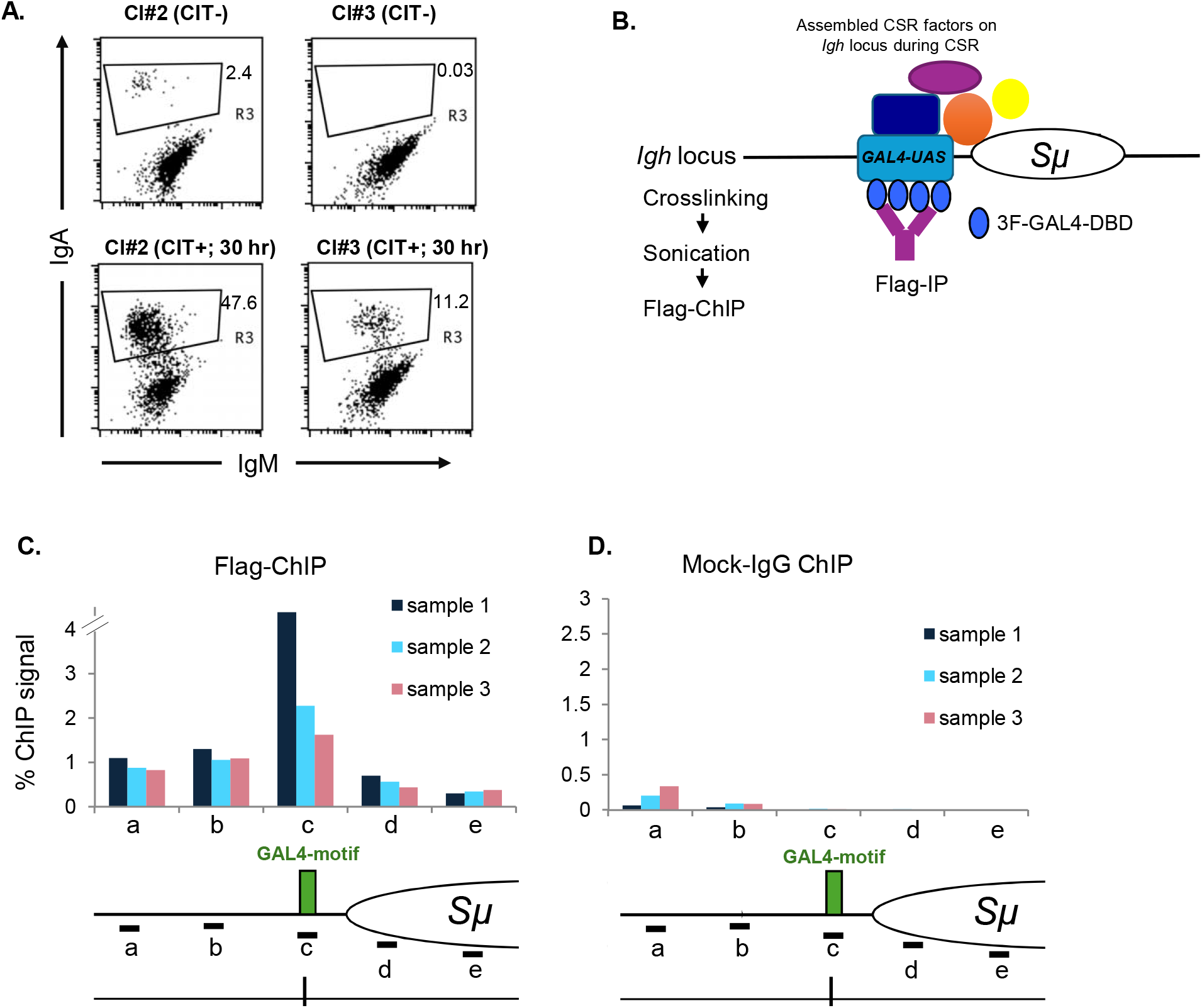
Clones inserted with *5X-GAL4-UAS* and stably overexpressing the 3F-GAL4-DBD undergo CSR. **(A)** Cl-2 and CL-3 were stimulated with CD40 ligand, IL-4 and TGF-β (CIT) with doses indiated previously [13]. After 30 hours of CIT stimulation, flow cytometry was performed to quantify the surface level expresion of IgM and IgA. IgA switching (CSR efficiency) in the Cl-2 and Cl-3 was 47.6% and 11.2 % respectively. **(B)** Schematic representation of 3F-GAL4-DBD binding to sonicated fragments of *5X-GAL4-US* inserted locus, enriched with CSR specific factors. The 3F-GAL4-DBD protein binds *5X-GAL4-UAS*, enabling the pulldown of additional factors assembled in its vicinity during CSR induction. **(C-D)** Flag-ChIP and IgG ChIPs were performed in *5X-GAL4-UAS* inserted CH12 clones stably overexpressing the 3F-GAL4-DBD. Three replicates were used for the ChIP analysis. The schematic of *IgH* locus is shown in the *5X-GALL4-UAS* clones indicating the position of primer pairs a, b, c, d, and e. Percentage of ChIP signal normalized to input values are indicated. IgG ChIP is used as a control for background levels of ChIP signal with same amount of chromatin input.

We next checked whether the 3F-GAL4-DBD protein can bind to the *Igh* locus. We performed the Flag Chromatin Immunoprecipitation (ChIP) assay using the triplicate samples of the Cl-2 (Figure 4B). Flag-ChIP revealed a significant enrichment of 3F-GAL4-DBD at the *Igh* locus compared to the mock IgG ChIP samples at all five locations examined. The Flag-ChIP signals were detected on five locations, indicated with primer pairs *a, b, c, d*, and *e* (Figure 4B). The significant enrichment of 3F-GAL4-DBD at five different loci suggested that 3F-GAL4-DBD is efficiently tethered to the *Sµ* region on the *Igh* locus (Figure 4B-C). These results confirm the successful establishment of the *5X-GAL4-UAS* cell line with 3F-GAL4-DBD expression, allowing the isolation CSR factor from the *Sµ* region during the CSR.

### Pulldown of CSR-specific factors from *5X-GAL4-UAS* inserted CH12 cells

We confirmed that *5X-GAL4-UAS* cells stably expressing 3F-GAL4-DBD exhibited normal tethering of the 3F-GAL4-DBD in the near vicinity of *Sµ* regions (Figure 4C-D). We next attempted to detect BRD2, a positive regulator of CSR enriched at *Sµ* during CSR. BRD2, a bromodomain protein that positively regulated CSR by promoting non-homologous end joining [13]. To detect BRD2 at the *Sµ* region, we performed 3F-GAL4-DBD ChIP after 24 hours of CIT induction in these clones under either scramble or *Brd2* knockdown conditions (Figure 5A, B).

**Figure 5.**
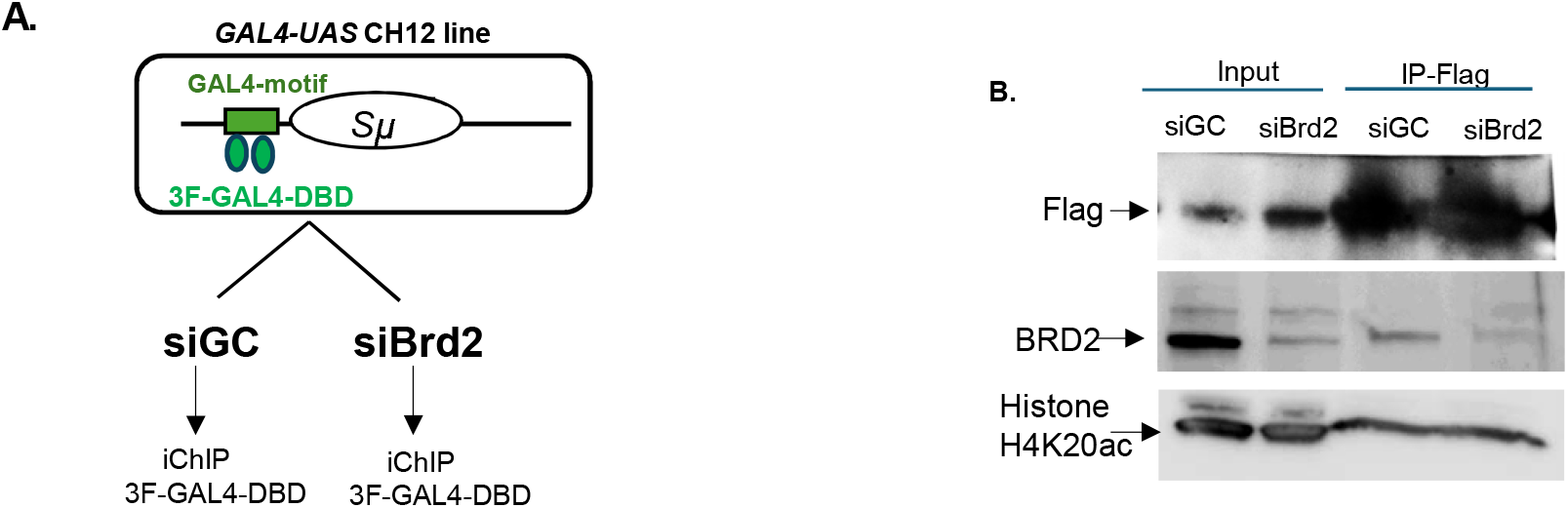
Pulldown of 3F-GAL4-DBDB binding proteins from the *5X-GAL4-UAS* sequence. **(A)** Schematic of CSR induction in *5X-GAL4-UAS* inserted CH12 cells in either *siControl* or *siBrd2* depleted cells. CSR was induced by CIT induction for 30 hours and nuclear lysate was prepared for Flag-IP. **(B)** The protein eluates from Flag-IP were subjected to Western blotting analysis against Flag, BRD2, and histone H4 lysine 20 acetylation. Input is 10% of total IPed sample. Flag-IP elute from *siBrd2* knockdown cells exhibited reduced signals of BRD2 on the *Igh* locus. BRD2 protein enrichment is used as a positive factor enriching to the *Ig* locus during the CSR, as described [13].

Upon performing the Flag-ChIP, we confirmed enrichment of the 3F-GAL4-DBD protein by immunoblotting in both scramble and *Brd2* knockdown cells (Figure 5B), indicating successful pulldown of BRD2 from the *Sµ* region. We observed higher BRD2 occupancy in the immunoprecipitates obtained from the scramble knockdown cells, while BRD2 signals were reduced in the immunoprecipitates obtained from *Brd2* knockdown cells (Figure 5B). These results demonstrate the successful assembly and isolation of CSR-specific protein complexes on the *Sµ* regions in *5X-GAL4-UAS* CH12 cells.

## Discussion

### Insertion of *5X-GAL4-UAS* into the *Igh* locus

Here we have successfully created a system to enrich for factors physically associating with *Sµ* region during CSR using the *GAL4* system and validated this tool by enriching for the chromatin organizer BRD2, a critical promoter of successful CSR. A similar approach could be applied with human B lymphoma cell lines harboring an intact *IG* locus, further allowing the identification of factors involved in safe execustion of CSR and suppression of the genetic rearrangements during the antibody isotype switching in human B cells.

The frequency of IgA class switching in Cl-2 and Cl-3 cells was 47.6% and 11.2%, respectively (Figure 4A), compared to the typical 10–20% class switch recombination (CSR) observed in normal CH12 cells after approximately 30 hours of stimulation [27]. Notably, Cl-2 exhibited a markedly higher CSR frequency, suggesting an enhanced responsiveness to CSR stimulation upon *GAL4-UAS* insertion into the *Sµ* region. The high frequency in IgA switching in Cl-2 suggests the *5X-GAL4-UAS* insertion positively impacts normal CSR. In future, comparison of protein complexes assembled at the *Sµ* region between Cl-2 to Cl-3 can define factors enhancing CSR frequency. Overall, the *5X-GAL4-UAS* insertion system offers a powerful platform to dissect the spatial and temporal dynamics of CSR factor recruitment, and mass spectrometry–based identification of eluted proteins will be essential for uncovering the key regulators of antibody class switching and maintaining genomic integrity.

## Acknowledgement

The research was supported by was supported by Takeda Science foundation scholarship and Kanehara foundations to SG. The author acknowledges Dr. Nasim A. Begum for helpful discussions. The author acknowledges Dr. Jacqueline H. Barlow for critical reading of the manuscript and constructive feedback.

## Author Contribution

SKG wrote the original draft, performed the experiment and analyzed the data.

## Methods

### Cell culture, transfection, selection against the resistance markers and CSR induction

CH12 cells were cultured as described [13]. Briefly 1 million cells were cultured and transfected using Lonza-nucleofector kit (Catalog#VPE-1001). The siRNA knockdown, control and *Brd2* knockdown was performed as described earlier [13].

### Construction of CRISPR-Cas9 plasmids and *5X-GAL4-UAS* targeting cassettes

The CRISPR/Cas9 plasmids targeting either *Sµ* or *Rosa-26* locus were constructed in GeneArt™ CRISPR Nuclease Vector with CD4 Enrichment Kit (Catalog# A21175, Invitrogen, Japan). Guide RNA sequences for CRISPR/Cas9 plasmids are described in Table 1. Homology arms 1 and 2 flanking about 1 kb upstream and downstream sequences of CRISCPR-Cas9 targeting site were amplified using genomic DNA (750 ng) obtained from wild type CH12 cells. PCR amplified fragments of homologous arms were cloned upstream and downstream of *5X-GAL4-UAS* sequences (120 bp) in the *pGL5* vector. Arm 1 was cloned upstream of *5X-GAL4-UAS* sequences in the *pGL5* vector using NotI/KpnI restriction sites. Arm 2 was cloned downstream of *5X-GAL4-UAS* sequences using NheI/BamHI restriction sites. The forward and reverse primer sequences for PCR amplification of homology arm 1 and arm 2 are described in Table 1. Homology arms were confirmed by DNA sequencing of the targeting construct. A linearized fragment was generated from the targeting cassette using NotI/BamHI sites, containing arm 1, *5X-GAL4-UAS* and arm 2. Transfection with 1μg of linearized fragment of the targeting cassette along with 1μg of CRISPR-Cas9 plasmid was performed into 1 million of CH12 cells. Cells were stained for CD4 and sorted using magnetic beads to enrich for CRISPR/Cas9-transfected cells, according to the manufacturer’s instructions (Catalogue#A21175, Invitrogen, Japan). From the sorted cells, single cell dilution was performed to obtain the single clones.

### Southern blotting for the confirmation of *5X-GAL4-UAS* insertions into the *Igh* locus

CD4 is co-expressed from the backbone of CRISPR/Cas9 plasmid and thus allows the sorting of the CRISPR-Cas9 transfected CH12 cells. CD4+ sorting was performed as per the manufacturer’s instruction to sort CH12 cells transfected with CRISPR/Cas9 plasmid targeting the 1 kb upstream of the *Sµ* region (*5X-GAL4-UAS* insertion site). After sorting transfected cells, genomic DNA was isolated, and PCR was performed to check integration of *5X-GAL4-UAS* plasmid. The primer sequences used to check *GAL4-UAS* integration were: forward primer 5’ GTAGGCCTGGACTTTGGGTCTCCC 5’ GCATCAGTACCTCAACTAATAC, respectively.

The PCR product was purified digested with EcoRI, giving 2 fragmetns from the wild type of amplicon 3 fragments in the amplification of *5X-GAL4-UAS* inserted clones. Ethidium bromide (0.5 µg/mL) staining was performed for 30 minutes to follwed by destaining for 30 minutes. Southern blotting and visualization of hybridized membranes was performed as described [13]. Probe sequences are described in table 1.

### Stable expresion of 3F-GAL4-DBD from *Rosa-26* locus

For site-directed insertion of *pEF1a-3FLAG-GAL4-DBD-IRES*-puromycin at the *Rosa-26* locus, the homology arms were constructed into zero-blunt vector (Invitrogen). The first and second homology arm flaking insertion site on *Rosa-26* locus were cloned using the primer sequences described into the Table 1. After cloning homology arms into the zero-blunt vector, the *pEF1a-3FLAG-GAL4-DBD-IRES*-puromycin fragment was PCR amplified using the forward primers sequence 5’ AGTCACGCGTGAGTAATTCATACAAAAGGAC and reverse primer sequence 5’ AGCTCATATGCCTCGTCTTGCAGTTCATTCA. Purified PCR product was cloned in middle of arm 1 and arm 2 of the zero-blunt vector using MluI and NdeI. CRISPR/Cas9 plasmids (1μg) and the targeting cassette (1μg) were electroported simulatenouly in the the *5X-GAL4-UAS* inserted clones. After 24 hr of transfection, puromycin was added to a concentration of 500μg/ml and cells were allowed for incubation for two days. The cells were cultured in puromycin for one week, with fresh medium containing the indicated doses of puromycin replaced every three days.

### Flag Immunoprecipitation, and Western blotting

5x10^7^ cells were Crosslinked at room temperature with 1% formaldehyde for 10 min. Crosslinking was stopped by adding Glycine solution (2 ml of 2M glycine in 30 ml for 5 min). Cells were washed twice with room temperature PBS. Nuclei were prepared with cell lysis kit (Thermo NE-PER 78833). The nuclei were sonicated (30 sec on, 60 sec rest for 9 cycles). Sonicated nuclei were spun at 13,500g for 5 minutes. Supernatant was collected and incubated with Flag-agarose beads (Sigma M2). IP reaction was incubated at 4 degree celsius overnight. IP-ed beads were washed three times (5 minutes each) with the Pierce IP lysis solution (Thermos#87787). IPed samples were eluted using 10 µl Flag peptide diluted in the 40 µl of PBS. For elution, beads and Flag peptide were kept on ice. Elutes were mixed with 40µl of 2X Sample buffer containing β-mercaptoethanol. The samples were boiled at 95 degree celsius for 5 minutes and analyzed by western blotting.

### Chromatin Immunoprecipitation and qRT-PCR

ChIP assay was performed as described [13]. Briefly, 10 million cells were crosslinked with 1% paraformaldehyde (Invitrogen) for 10 min followed by formaldehyde-quenching with glycine treatment [13]. The crosslinked cells were lysed, nuclei were isolated and sonicated to obtain average fragment size of chromatin about 500-700 bp. The Flag-IP and mock IgG IP was performed using the M2-Flag antibody (Sigma) and a control Ig Isotype antibody (Sigma). The IPed chromatin was washed and DNA-was purified using the Zymo research ChIP-DNA cleaning kit (Catalogue#D5205). The binding of 3F-GAL4-DBD to the S region was assessed at 5 loci by qPCR analysis using the primer pairs binding to regions a, b, c, d, e. Primer sequences are described in Table 1.

